# Imaging object-scene integration in visible and invisible natural scenes

**DOI:** 10.1101/116111

**Authors:** Nathan Faivre, Julien Dubois, Naama Schwartz, Liad Mudrik

## Abstract

Integrating objects with their context is a key step in the interpretation of complex visual scenes. Humans can do this very quickly, yet the brain mechanisms that mediate this ability are not yet understood. Here, we used functional Magnetic Resonance Imaging (fMRI) to measure brain activity while participants viewed visual scenes depicting a person performing an action with an object that was either congruent or incongruent with the scene. Univariate and multivariate analyses revealed different activity for congruent compared to incongruent scenes in the lateral occipital complex, inferior temporal cortex, parahippocampal cortex, and prefrontal cortex, in line with existing models of scene processing. Importantly, and in contrast to previous studies, these activations could not be explained by task-induced conflicts. A secondary goal of this study was to examine whether object-context integration could occur in the absence of awareness, by comparing brain activity elicited by congruent vs. incongruent scenes that were suppressed from awareness using visual masking. We found no evidence for brain activity differentiating between congruent and incongruent invisible scenes. Overall, our results provide novel support for the roles of PHC and PFC in conscious object-context integration which cannot be explained by either low-level differences or task demands. Yet they further suggest that activity in these regions is decreased by visual masking to the point of becoming undetectable with our fMRI protocol.

## 1. Introduction

A very short glimpse of a visual scene often suffices to identify objects, and understand their relations with one another, as well as with the context in which they appear. The co-occurrence of objects within specific scenes or contexts (Bar, 2004) gives rise to expectations about their relations. When these expectations are violated (Biederman, Mezzanotte, & Rabinowitz, 1982), object and scene processing are impaired - both with respect to speed (e.g., Davenport & Potter, 2004; Palmer, 1975; Rieger, Kochy, Schalk, Gruschow, & Heinze, 2008) and to accuracy (e.g., Biederman, Rabinowitz, Glass, & Stacy, 1974; Boyce, Pollatsek, & Rayner, 1989; Underwood, 2005), suggesting that contextual expectation may have an important role in scene and object processing (Bar, 2004; Mudrik, Lamy, & Deouell, 2010; Oliva & Torralba, 2007; though see Henderson & Hollingworth, 1999).

Yet despite the proclaimed role of object-context integration in scene comprehension, its underlying mechanisms are still unclear. At the neural level, an interplay between frontal and temporal visual areas has been suggested (Bar, 2003, 2004; see also Trapp & Bar, 2015), so that after identifying the scene’s gist, high-level contextual expectations about scene-congruent objects are compared with upcoming visual information about objects’ features, until a match is found and the objects are identified (for electrophysiological support, see Dyck & Brodeur, 2015; Mudrik et al., 2010; Mudrik, Lamy, Shalgi, & Deouell, 2014; Võ & Wolfe, 2013; though see Ganis & Kutas, 2003; Demiral et al., 2012). However, these suggestions are mostly based on studies that did not directly examine the processing of objects in scenes, but rather used other ways to probe contextual processing (e.g., comparing objects that evoke strong vs. weak contextual associations (Kveraga et al., 2011), or manipulating the relations between two isolated objects (Gronau, Neta, & Bar, 2008; Kaiser, Stein, & Peelen, 2014)). Critically, the few papers that did focus on objects within real life scenes (Jenkins, Yang, Goh, Hong, & Park, 2010; Kirk, 2008; Rémy, Vayssière, Saint-Aubert, Barbeau, & Fabre-Thorpe, 2013) report conflicting findings and interpretations about the role of frontotemporal regions - more specifically the prefrontal cortex and the medial temporal lobe (MTL) - for object-context integration. The prefrontal cortex was repeatedly implicated in semantic processing of different types (e.g., Gold et al., 2006; Gronau et al., 2008; Kveraga et al., 2011; van Kesteren, Rijpkema, Ruiter, & Fernández, 2010), even when directly manipulating object-scene relations (Rémy et al., 2013). Yet it was suggested that this may reflect task-induced conflict rather than object-context integration per se (Rémy et al., 2013). Likewise, the literature is divided about the role of the MTL in object-context integration, especially regarding the parahippocampal cortex (PHC; for review of findings, see Malcolm, Groen, & Baker, 2016). Some claim that the PHC processes contextual associations (Aminoff, Kveraga, & Bar, 2013; Bar & Aminoff, 2003; Montaldi et al., 1998; Rémy et al., 2013; see also Goh et al., 2004, who reported PHC and hippocampal activations for contextual binding between objects and scenes, and Stansbury et al. (2013) who gave an account of PHC activity in terms of statistical learning of object co-occurrences). Others argue that it processes spatial layouts (Epstein, 2008; Epstein & Ward, 2009) or representations of three-dimensional local spaces, even of a single object (Mullally & Maguire, 2011), irrespective of contextual associations (see also Howard, Kumaran, Ólafsdóttir, & Spiers, 2011).

The cognitive characteristics of object-context integration are no better agreed upon than its neural mechanisms. For instance, the necessary conditions for such integrative processes are still under dispute. One aspect of this dispute, which is at the focus of the current research, concerns the role of conscious perception in object-context integration. Using behavioral measures, two studies suggested that integration can occur even when subjects are unaware of both objects and the scenes in which they appear (Mudrik, Breska, Lamy, & Deouell, 2011; Mudrik & Koch, 2013; see also Stein, Kaiser, & Peelen, 2015, for prioritized access to awareness of interacting vs. non-interacting objects). This result is in line with the *unconscious binding hypothesis* (Lin & He, 2009), according to which the brain can associate, group or bind certain features in invisible scenes, especially when these features are dominant (for a discussion of conscious vs. unconscious integration, see Mudrik, Faivre, & Koch, 2014). However, a recent attempt to replicate these findings has failed (Moors, Boelens, van Overwalle, & Wagemans, 2016; Kataev & Mudrik, under review). The absence of unconscious object-context integration would be in line with theories that tie integration with consciousness (Global Workspace Theory, Dehaene & Changeux, 2011; Dehaene & Naccache, 2001; Integrated Information Theory, Tononi, Boly, Massimini, & Koch, 2016; Tononi & Edelman, 1998). While the jury is still out on this question, our study aimed at measuring the brain activity mediating unconscious object-context integration - if indeed such integration is possible in the absence of awareness.

The goals of the current study were thus twofold; at the neural level, we aimed at identifying the neural substrates of object-context integration, and specifically at testing whether frontal activations indeed reflect contextual processing rather than task-related conflicts (Rémy et al., 2013). At the cognitive level, we looked for evidence of unconscious object-context integration while carefully controlling visibility, and aimed at identifying the neural substrates of unconscious integration. Subjects were thus scanned as they were presented with masked visual scenes depicting a person performing an action with a congruent (e.g., a man drinking from a bottle) or an incongruent (e.g., a man drinking from a flashlight) object (see Figure 1). The experiment had two conditions: one in which scenes were clearly visible (visible condition), and one in which they were not (invisible condition). Stimulus visibility was manipulated by changing masking parameters while keeping stimulus energy constant. Participants rated stimulus visibility on each trial; they did not perform any object-context congruency or object-identification judgments, to ensure that the measured brain activations could be attributed to object-context integration per se, and not to task-induced conflict (Rémy et al., 2013).

**Figure 1.**
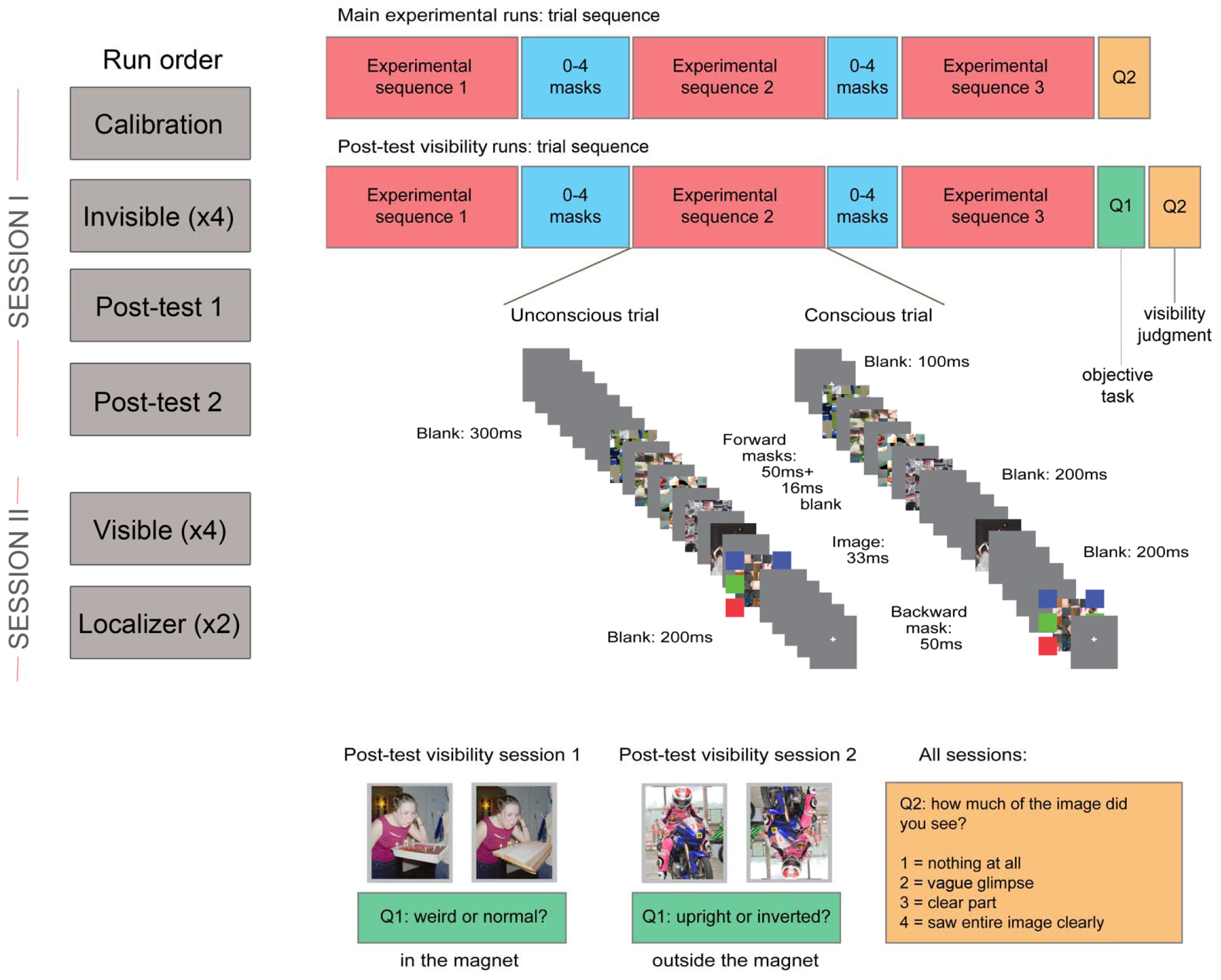
Experimental procedure. The left column depicts the different runs and their order (calibration, invisible session, visibility tests, visible session and localizer). On the right, the experimental sequence in the main runs (top) and the two post-test visibility runs (bottom). In all trials, one experimental sequence was repeated three times, separated by 0-4 masks to avoid temporal predictability of the target image. The experimental trials ended with only one question about target visibility. In the post-test trials, this subjective visibility question was preceded by an objective question (about target congruency/orientation). Each experimental sequence included one presentation of the target scene, which was either congruent or incongruent, and five masks which either immediately followed and preceded the scene (invisible condition) or were separated from it by blanks (visible condition). Thus, throughout each trial, the target scene was presented three times.

## 2. Methods

### 2.1 Participants

Eighteen participants (eight females, mean age = 25.1 years, SD = 4.32 years) from the student population of the California Institute of Technology took part in this study for payment ($50 per hour). All participants were right-handed, with normal or corrected-to-normal vision, and no psychiatric or neurological history. The experiment was approved by the Institutional Review Board committee of the California Institute of Technology, and informed consent was obtained after the experimental procedures were explained to the subjects. Two additional participants had too few invisible trials in the invisible condition (less than 70% of trials), and were excluded from the analysis.

### 2.2 Apparatus and stimuli

Stimuli were back-projected onto a screen that was visible to subjects via a mirror attached to the head coil, using a video projector (refresh rate 60 Hz, resolution 1280 x 1024). Stimuli were controlled from a PC computer running Matlab with the Psychophysics Toolbox version 3 (Brainard, 1997; Kleiner et al., 2007; Pelli, 1997). Responses were collected with two fiber optic response devices (Current Designs, Philadelphia, USA). Target images were 6.38° by 4.63° (369 x 268 pixels) color pictures of a person performing an action with an object. In the congruent condition, the object was congruent with the performed action (e.g., a woman drinking from a cup), while in the incongruent condition, it was not (e.g., a woman drinking from a plant; see Mudrik, Deouell, & Lamy, 2011; Mudrik & Koch, 2013 for details). In both types of images, the critical object was pasted onto the scene. Low-level differences in saliency, chromaticity, and spatial frequency were controlled for during the creation of the stimulus set (Mudrik et al., 2010), and tested using an objective perceptual model (Neumann & Gegenfurtner, 2006). Visual masks were generated from a different set of scenes, by cutting each scene image into 5 x 6 tiles and then randomly shuffling the tiles.

### 2.3 Procedure

The experiment was run over two separate one-hour scanning sessions. During the anatomical scan at the beginning of the first session, subjects performed a simple discrimination task designed to individually adjust the contrasts of the masks and targets, thus ensuring a comparable depth of suppression across subjects in the invisible condition. 72 images (half congruent, half incongruent; all different from the ones used in the main experiment) were presented, either upright or inverted (pseudo-randomly intermixed, with the constraint that the same image orientation was never presented in four consecutive trials). Subjects indicated for each trial whether the image was upright or inverted, and rated its visibility subjectively using the Perceptual Awareness Scale (PAS; Ramsøy & Overgaard, 2004), where 1 signifies ‘‘I didn’t see anything,’’ 2 stands for ‘‘I had a vague perception of something,’’ 3 represents ‘‘I saw a clear part of the image,’’ and 4 is ‘‘I saw the entire image clearly.’’ Subjects were instructed to guess the orientation if they did not see the image. Initial mask (Michelson) contrast was 0.85, and initial prime contrast was 0.7. Following correct responses (i.e., correct classification of the image orientation as upright or inverted), mask contrast was increased by 0.05, and following incorrect responses, it was decreased by 0.05 (i.e., 1-up, 1-down staircase procedure, Levitt, 1971). If mask contrast reached 1, target contrast was decreased by steps of 0.05, stopping at the minimum allowable contrast of 0.15. In the main experiment, mask contrast was then set to the second highest level reached during the calibration session (or, if mask contrast reached 1, it was set to 1) and target contrast was set to the second lowest level reached. Mask contrast reached 1 for all subjects. Average target contrast was 0.39 ± 0.05 (here and elsewhere, ± denotes 95% confidence interval). The same contrasts were used in both visible and invisible conditions in the main experiment, so stimulus energy entering the system would be matched.

The invisible condition followed the calibration in the first session. The visible condition was conducted in the second session, so the results in the invisible condition would not be biased by previous conscious exposure. The invisible and visible conditions were each divided into four runs of 90 trials, of which 72 contained either congruent or incongruent target images, and 18 had no stimuli (i.e., “catch trials”), serving as baseline. The order of congruent and incongruent trials within a run was optimized using a genetic algorithm (Wager & Nichols, 2003) with the constraint that a run could not start with a baseline trial, and that two baseline trials could not occur in succession. Each trial started with a fixation cross presented for 200 ms, followed by three repeats of a sequence of target images and masks, and then a judgment of image visibility using the PAS. The sequence started with two forward masks (each presented for 50 ms, with a 17 ms blank interval), followed by the target image (33 ms), and two backward masks (50 ms each, 17 ms blank interval). The only difference between the invisible condition (first session) and the visible condition (second session) lied in the duration of the intervals immediately preceding and immediately following the target image: a 17ms gap was used in the invisible session, while a 50 to 200 ms gap (randomly selected on each trial from a uniform distribution) was used in the visible session. To equate the overall energy of a trial across conditions, the final fixation in the sequence lasted between 100 and 400 ms in the invisible session. A random number of masks (0-4) were presented between repetitions of the sequence in each trial, to minimize predictability of the onset of the target image (Figure 1). A random inter-trial interval (uniform distribution between 1 and 3s) was enforced between trials, so that on average a whole trial lasted 4.5 s.

At the end of the invisible session, two objective performance tasks were run in the scanner: 1) a congruency task, in which subjects were asked to determine if a scene was congruent or not and 2) an orientation task, in which half the images were upright and half were inverted, and subjects were asked to determine their orientation. In both tasks, the same trial structure as the one used in the main experiment was used; subjects were instructed to guess if they did not know the answer. Subjects also rated image visibility using the PAS, after each trial.

At the end of the visible session, subjects participated in two runs of a block design paradigm to localize brain regions that respond differentially to congruent and incongruent scenes. Each consisted of 18 blocks of 12 images, which were either all congruent or all incongruent scenes. Blocks started with a 5-7 s fixation cross. Then, the 12 images were presented successively for 830 ms each, with a 190 ms blank between images. Subjects had to detect when an image was repeated, which occurred once per block (1-back task).

### 2.4 Behavioral data analysis

All analyses were performed with R (2016), using the BayesFactor (Morey & Rouder 2015), and ggplot2 (Wickham, 2009) packages.

### 2.5 MRI data acquisition and preprocessing

All images were acquired using a 3 Tesla whole-body MRI system (Magnetom Tim Trio, Siemens Medical Solutions) with a 32-channel head receive array, at the Caltech Brain Imaging Center. Functional blood oxygen level dependent (BOLD) images were acquired with T2*- weighted gradient-echo echo-planar imaging (EPI) (TR/TE = 2500/30 ms, flip angle = 80°, 3mm isotropic voxels and 46 slices acquired in an interleaved fashion, covering the whole brain). Anatomical reference images were acquired using a high-resolution T1-weighted sequence (MPRAGE, TR/TE/TI = 1500/2.74/800 ms, flip angle = 10°, 1mm isotropic voxels).

The functional images were processed using the SPM8 toolbox (Wellcome Department of Cognitive Neurology, London, UK) for Matlab. The three first volumes of each run were discarded to eliminate nonequilibrium effects of magnetization. Preprocessing steps included temporal high-pass filtering (1/128 Hz), rigid-body motion correction and slice-timing correction (middle reference slice). Functional images were co-registered to the subject’s own T1-weighted anatomical image. The T1-weighted anatomical image was segmented into gray matter, white matter and cerebrospinal fluid, and nonlinearly registered to the standard Montreal Neurological Institute space (MNI152). The same spatial normalization parameters were applied to the functional images, followed by spatial smoothing (using a Gaussian kernel with 12 mm full-width at half maximum) for group analysis. Scans with large signal variations (i.e., more than 1.5% difference from the mean global intensity) or scans with more than 0.5 mm/TR scan-to-scan motion were repaired by interpolating the nearest non-repaired scans using the ArtRepair toolbox (Mazaika, Whitfield, & Cooper, 2005).

### 2.6 fMRI analysis

Statistical analyses relied on the classical general linear model (GLM) framework. For univariate analyses, models included one regressor for congruent, one regressor for incongruent, and one regressor for baseline trials in each run. Each regressor consisted in delta functions corresponding to trial onset times, convolved with the double gamma canonical hemodynamic response function (HRF). Time and dispersion derivatives were added to account for variability of the HRF across brain areas and subjects. Motion parameters from the rigid-body realignment were added as covariates of no interest (6 regressors). Individual-level analyses investigated the contrast between the congruent vs. the incongruent condition. Group-level statistics were derived by submitting individual contrasts to a one sample two-tailed t-test. We adopted a cluster-level thresholding with p < 0.05 after FWE correction, or uncorrected threshold with p < 0.001 and a minimal cluster extent of 10 voxels. Note that our results did not resist correction for multiple comparisons after threshold free cluster enhancement (TFCE; Smith & Nichols, 2009). The functional localizer followed the same preprocessing and analysis steps as the main experimental runs. We defined regions of interest (ROIs) based on the results from the functional localizer: we centered spheres of 12 mm radius on the peak activity of each cluster of more than five contiguous voxels in the group-level SPM (voxel-wise threshold p < 0.001, uncorrected). For multivariate analyses, each run was subdivided into four mini-runs, each containing nine congruent and nine incongruent trials. One regressor was defined for congruent and incongruent trials in each mini-run. No time or dispersion derivative was used for these models, and no spatial smoothing was applied. The individual beta estimates of each mini-run were used for classification. Beta estimates of each voxel within the ROIs defined above were extracted and used to train a linear Support Vector Machine (using the libsvm toolbox for Matlab, http://www.csie.ntu.edu.tw/cjlin/libsvm to classify mini-runs into those with congruent and incongruent object-context relations. A leave-one-run-out cross-validation scheme was used. Statistical significance of group-level classification performances was assessed using Bayes Factor and permutation-based information prevalence inference to compare global vs. majority null hypotheses (Allefeld, Görgen, & Haynes, 2016).

## 3. Results

### 3.1 Behavioral data

In the visible condition, subjects’ visibility ratings indicated that target images were partly or clearly perceived (visibility 1: 3.5% ± 3.0%; visibility 2: 18.5% ± 7.5%; visibility 3: 37.8% ± 5.4%; visibility 4: 45.4% ± 9.4%). Only trials with visibility ratings of 3 or 4 were kept for further analyses. By contrast, in the invisible condition, subjective visibility of target images was dramatically reduced due to masking (visibility 1: 56.6% ± 12.2%; visibility 2: 31.2% ± 7.1%; visibility 3: 14.3% ± 7.8%; visibility 4: 0.1% ± 0.04%) (Figure 2a). Only trials with visibility ratings of 1 or 2 were kept for further analyses (for a similar approach, see Melloni, Schwiedrzik, Müller, Rodriguez, & Singer, 2011). Objective performance for discriminating scene congruency in corresponding visibility ratings in the invisible condition did not significantly differ from chance-level (51.6 % ± 2.5%, t(15) = 1.24, p = 0.23, BF = 0.49, indicating that H0 was two times more likely than H1) (Figure 2b). However, objective performance for discriminating upright vs. inverted scenes was slightly above chance-level (56.0 % ± 4.4%, t(16) = 2.64, p = 0.02, BF = 3.3, indicating that H1 was around three times more likely than H0; regrettably, two subjects did not complete the objective tasks due to technical issues). This suggests some level of partial awareness, in which some participants could discriminate low-level properties of the natural scene such as its vertical orientation, but not its semantic content (Gelbard-Sagiv, Faivre, Mudrik, & Koch, 2016; Kouider & Dupoux, 2004). In all following results, unconscious processing is therefore defined with respect to scene congruency, and not to the target images themselves of which subjects might have been partially aware, at least in some trials.

**Figure 2.**
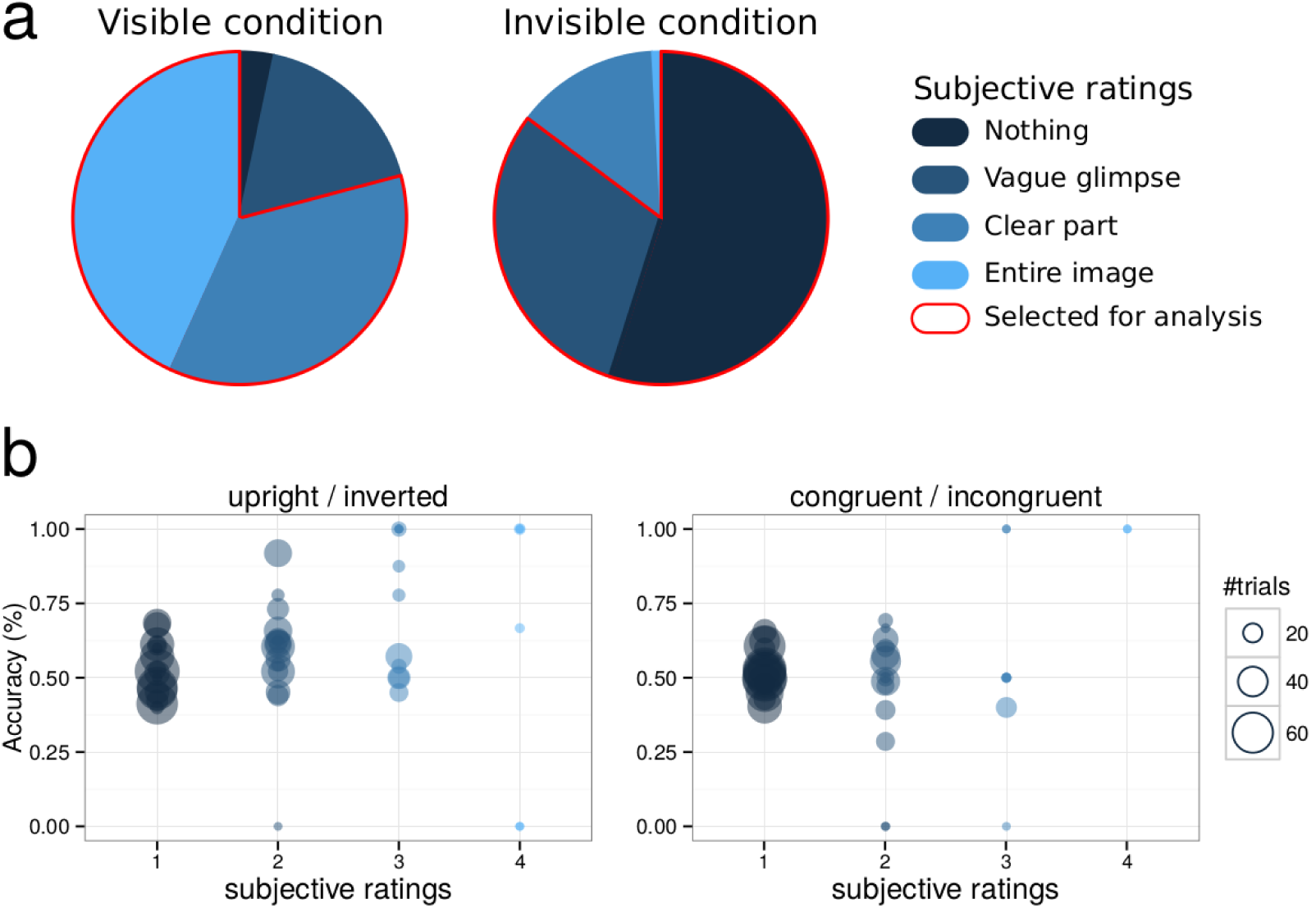
Behavioral results. 2a. Average distributions of subjective ratings in the visible and invisible conditions. In the visible condition, only trials in which participants reported seeing a clear part or the entire image were selected for analysis. In the invisible condition, these trials were excluded, while those in which nothing or a vague glimpse was perceived were kept. 2b. Objective performance for discriminating upright/inverted scenes (left panel) and congruent/incongruent scenes (right panel) in the invisible condition. Each circle represents individual performance in one category of subjective visibility. The size of each circle represents the number of trials from which individual performance was computed.

### 3.2 Imaging data

#### 3.2.1. Univariate analyses

The set of brain regions involved in the processing of scene congruency was identified with a functional localizer contrasting activity elicited by congruent vs. incongruent blocks of target images (no masking, see Methods). At the group level, this contrast revealed a set of regions in which activity was weaker for congruent than incongruent images. Significant activations (for cluster-level thresholding with p < 0.05 after FWE correction, or uncorrected threshold with p < 0.001 and a minimal cluster extent of 10 voxels) in the temporal lobe comprised the right inferior temporal and fusiform gyri, the left middle temporal gyrus, and the right parahippocampal gyrus including the parahippocampal place area. In the parietal lobe, activations were detected in the bilateral cunei, the right parietal lobule, and the left paracentral lobule. In the frontal lobe, significant clusters were observed in the bilateral inferior frontal gyri, right middle frontal gyrus, left superior medial frontal gyrus, and left cingulate gyrus.

We then used results from the localizer as a mask when contrasting activity elicited by congruent vs. incongruent target images during the visible condition in the main experimental runs (see Figure 3 and Table 2). Significant activations were found bilaterally in the fusiform, cingulate, middle and inferior frontal gyri, inferior parietal lobules, precunei, insular cortices, and caudate bodies; as well as activations in the left hemisphere, including the left inferior and superior temporal gyri, and left medial frontal gyrus (see Table 2).

**Figure 3.**
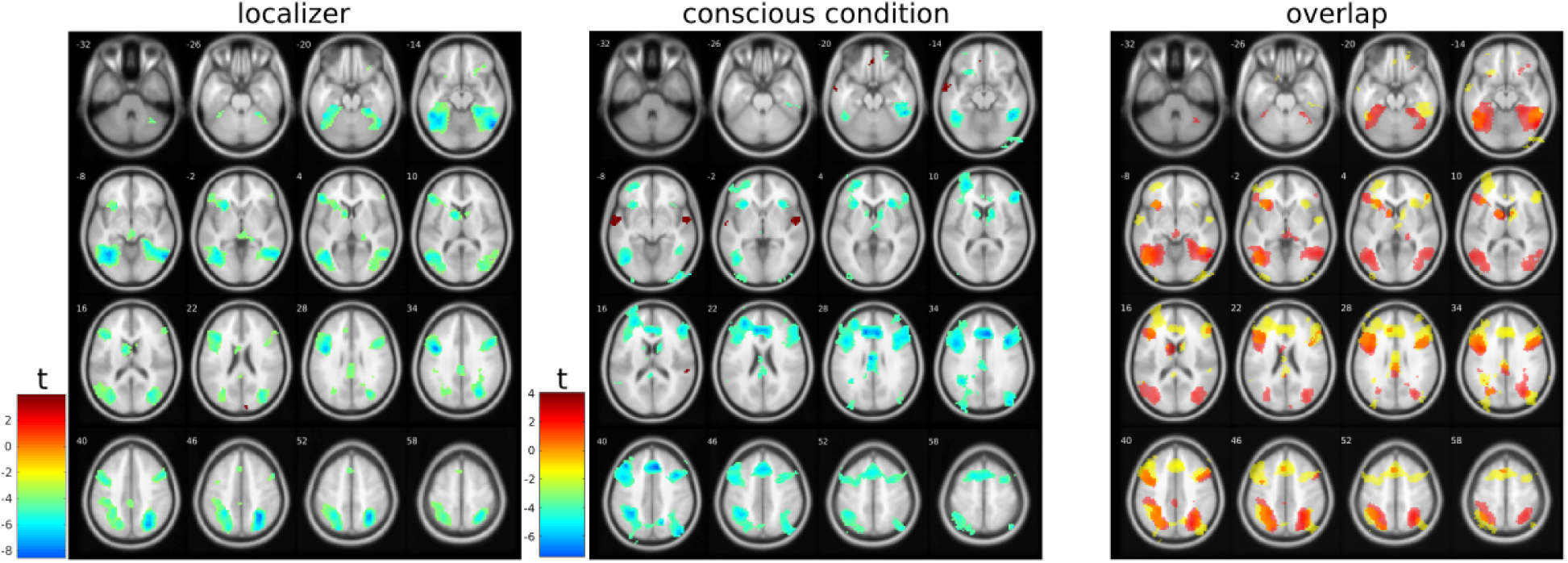
T-maps for the comparison of congruent vs. incongruent images during the functional localizer and visible condition (p < 0.001, uncorrected; see tables for corrected p-values). The right panel shows the overlap of t-values between the localizer (in red) and visible condition (in yellow). Activations in regions which were not identified with the localizer run included the following areas: in the right hemisphere, the inferior occipital, medial and superior frontal, and orbital gyri. In the left hemisphere – the superior and middle occipital, parahippocampal, fusiform, inferior temporal, supramarginal, and middle frontal gyri, as well as the inferior parietal lobule, posterior cingulate cortex, and caudate body.

**Table 1.** Cluster extent (CE), statistical values, and MNI stereotaxic brain atlas coordinates for the brain regions more activated by incongruent vs. congruent target images during the functional localizer.

| Cluster size | T-value (peak) | P-value | correction | x | y | z | Anatomical region | Broadmann area |
| --- | --- | --- | --- | --- | --- | --- | --- | --- |
| **2365** | 8.47 | < 0.001 | FWE | 57 | -61 | -11 | Right Inferior Temporal Gyrus | BA 37 |
|  | 8.27 |  | FWE | 30 | -55 | 49 | Right Precuneus | BA 7 |
|  | 7.41 |  | FWE | 27 | -70 | 40 | Right Superior Parietal Lobule | BA 7 |
| **1215** | 7.81 | < 0.001 | FWE | -42 | 2 | 34 | Left Inferior Frontal Gyrus | BA 9 |
|  | 6.28 |  | FWE | -27 | 17 | -2 | Left Claustrum |  |
| **2538** | 7.72 | < 0.001 | FWE | -33 | -58 | -11 | Left Cerebellum |  |
|  | 7.11 |  | FWE | -48 | -49 | -14 | Left Sub-Gyral | BA 37 |
|  | 7.03 |  | FWE | -36 | -67 | -5 | Left Middle Occipital Gyrus | BA 37 |
| **347** | 5.74 | 0.001 | FWE | 48 | 14 | 34 | Right Middle Frontal Gyrus | BA 9 |
|  | 4.51 |  | FWE | 57 | 35 | 19 | Right Middle Frontal Gyrus | BA 46 |
| **143** | 4.98 | 0.033 | FWE | 0 | -31 | 31 | Left Cingulate Gyrus | BA 23 |
|  | 3.88 | < 0.001 | unc | 0 | -34 | 46 | Left Paracentral Lobule | BA 31 |
| **55** | 4.41 | < 0.001 | unc | -3 | 20 | 52 | Left Medial Frontal Gyrus | BA 8 |
|  | 3.77 | < 0.001 | unc | -12 | 23 | 67 | Left Superior Frontal Gyrus | BA 6 |
| **45** | 4.26 | < 0.001 | unc | 24 | 20 | -14 | Right Inferior Frontal Gyrus | BA 47 |
|  | 4.22 | < 0.001 | unc | 21 | 35 | -17 | Right Middle Frontal Gyrus | BA 11 |
|  | 4.02 | < 0.001 | unc | 39 | 32 | -14 | Right Inferior Frontal Gyrus | BA 47 |
| **15** | 4.14 | < 0.001 | unc | 9 | 11 | 10 | Right Caudate | Caudate Body |
| **16** | 4.11 | < 0.001 | unc | -9 | 35 | 28 | Left Anterior Cingulate | BA 32 |
| **10** | 4 | < 0.001 | unc | -9 | -73 | -35 | Left Cerebellum |  |
| **19** | 3.93 | < 0.001 | unc | 51 | 38 | -2 | Right Middle Frontal Gyrus | BA 47 |

**Table 2:**
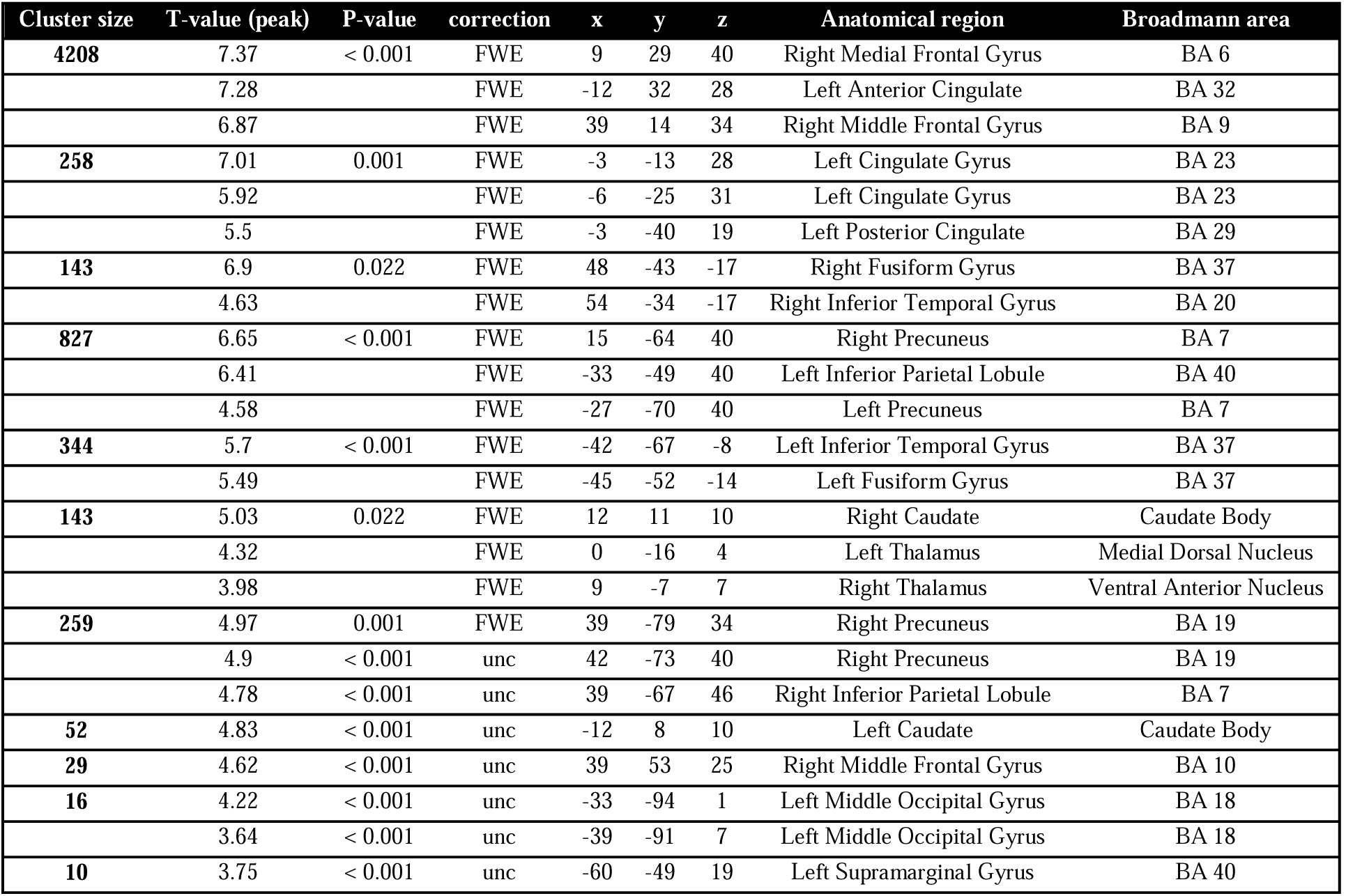
Cluster extent (CE), statistical values, and MNI stereotaxic brain atlas coordinates for the brain regions more activated by incongruent vs. congruent target images during the main task in the visible condition.

When contrasting activity elicited by congruent vs. incongruent target images in the invisible condition, no significant result was found with the same uncorrected threshold of p < 0.001. We verified that the images did elicit a response in visual cortices irrespective of their congruency, by contrasting both congruent and incongruent trials with the baseline condition in which no target image was shown (see figure SI1). Thus, masking was not so strong as to completely prevent low-level visual processing of the target images.

#### 3.2.2. Multivariate analyses

We then investigated whether the activity patterns extracted from spheres centered on the clusters identified in the functional localizer conveyed additional information regarding scene congruency. In the visible condition, we could decode scene congruency from the spheres located in the right fusiform gyrus, right superior parietal lobule, middle and inferior frontal gyrus, left precuneus, left superior medial and inferior frontal gyrus, and left anterior cingulate cortex. (Figure 4, Table 3). In the invisible condition, on the other hand, we could not decode congruency above chance in any of the spherical ROIs, with a statistical significance threshold of p < 0.05 for the global and majority null hypotheses (see Methods). Bayesian analyses revealed that our results in the invisible condition were inconclusive: evidence was lacking in our data to support the null hypothesis (i.e., log(1/3) < log(BF) > log(3), see Figure 4).

**Figure 4.**
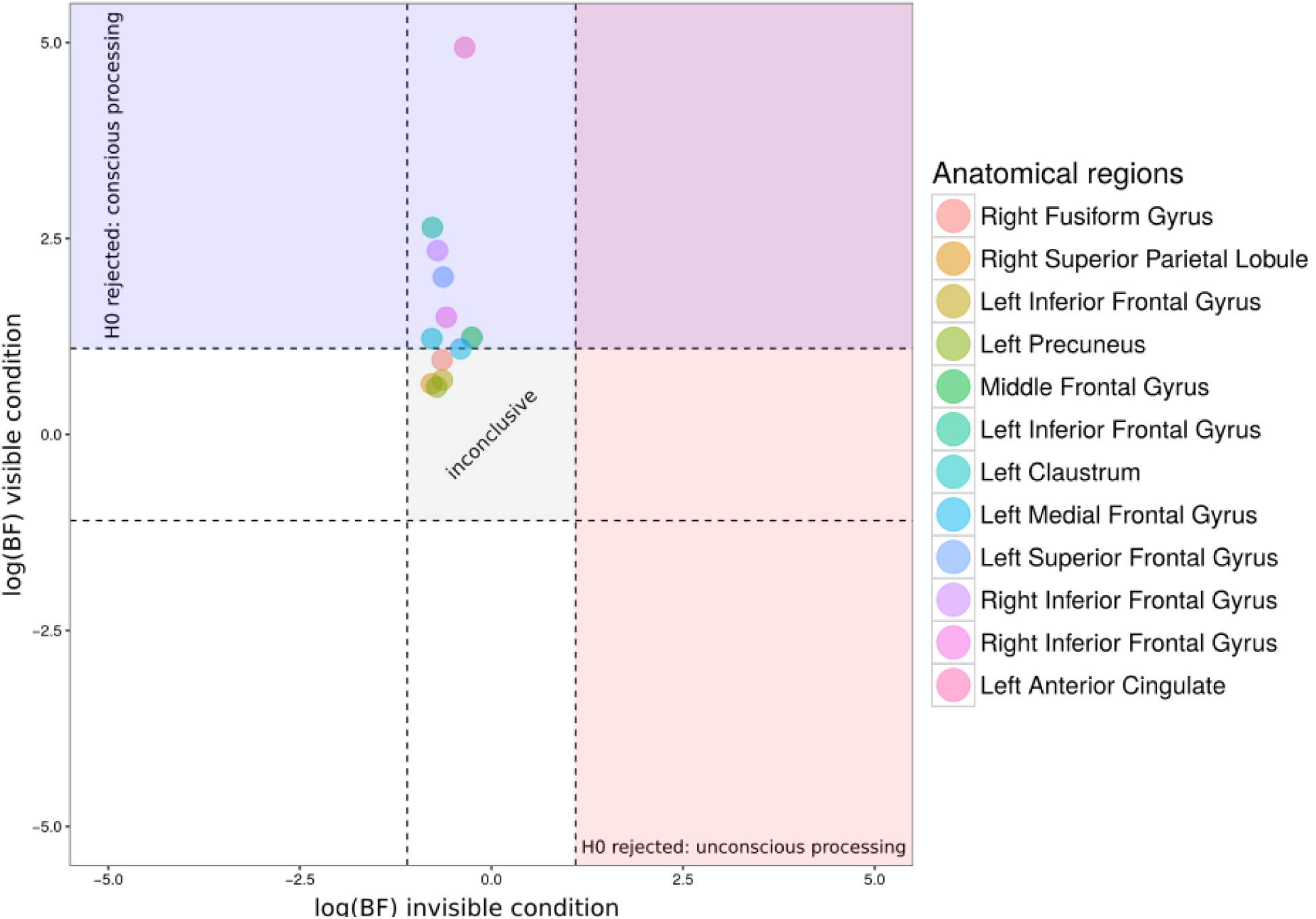
Logarithm of Bayes factor corresponding to decoding performances in the visible (y-axis) and invisible (x-axis) conditions, in the spherical ROIs where significant decoding was found in the visible condition. Dashed lines represent values below which the null hypothesis is favored (horizontal and vertical intercepts of log(⅓)), and above which the null hypothesis can be rejected (horizontal and vertical intercepts of log(3)). Note that all results in the invisible condition lay in the “gray zone” where the null hypothesis can neither be supported or rejected, suggesting inconclusive data; while most areas are above the dashed line in the visible condition, where the null hypothesis is rejected.

**Table 3:**
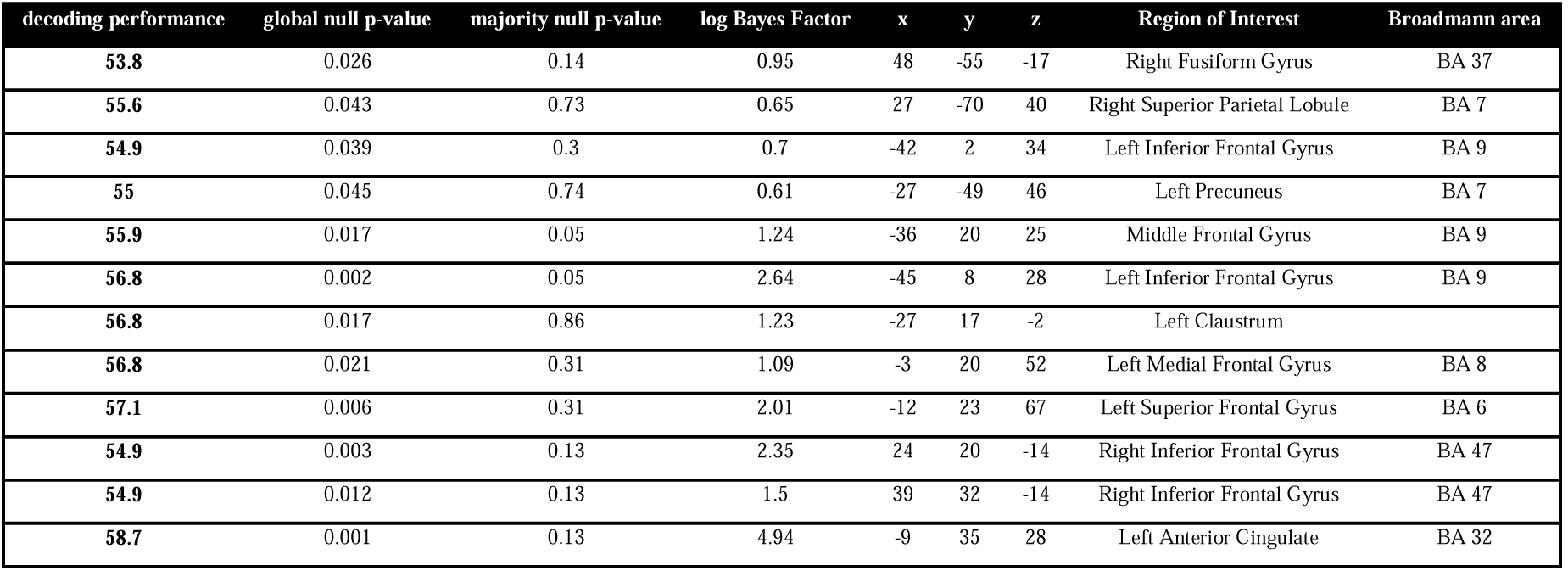
Average decoding performance, p-values for the global and majority null hypotheses (permutation-based information prevalence inference), logarithm of Bayes factors in the visible condition in the main task, and MNI stereotaxic brain atlas coordinates for the spheres centered on brain regions defined in the functional localizer. The majority null hypothesis was rejected at a threshold p<0.05 only in the left middle and inferior frontal gyri, suggesting that decoding was possible in a majority of participants in these brain regions.

## 4. Discussion

The current study aimed at determining the neural substrates of object-context integration that are recruited irrespective of a specific task or instruction, and assessing the existence of object-context integration even when subjects are unaware of the presented scenes. While for the first question an extensive network of areas engaged in object-context integration was identified − overcoming possible previous confounds induced by task demands or low-level features of the stimuli - the answer to the second question was negative, as we were unable to detect any neural activity which differentiated between congruent and incongruent scenes in the invisible condition. We discuss both of these findings below.

### 4.1. Neural correlates of object-context integration

In line with previous studies, several areas were found when contrasting activations induced by congruent vs. incongruent images. A prominent group of visual areas were identified: the inferior temporal gyrus, suggested by Bar (Bar, 2004; see also Trapp & Bar, 2015) as the locus of contextually-guided object recognition, and the fusiform gyrus, previously implicated also in associative processing (e.g., Hawco & Lepage, 2014). The fusiform gyrus comprises the lateral occipital complex (LOC), previously reported to differentiate between semantically related and unrelated objects (e.g., Kim & Biederman, 2010; see also Gronau et al., 2008). Interestingly, in a recent study which also presented congruent and incongruent scenes (Rémy et al., 2013), the LOC did not show stronger activity for incongruent images but only responded to animal vs. non-animal objects, irrespective of the scene in which they appeared. Possibly, this discrepancy could be explained by a difference in stimuli. All of our scenes portray a person performing an action with an object, so that object identity is strongly constrained by the scene (the number of possible objects which are congruent with the scene is very small; e.g., a person can only drink from a limited set of objects). In most previous studies, on the other hand, including that of Remy and colleagues, the scene is a location (e.g., a street, the beach, a living room), in which many objects could potentially appear. Potentially then, the stronger manipulation of congruency induced by our stimuli elicited wider activations.

Our results also support the role of the parahippocampal cortex (PHC) in processing contextual associations (Aminoff et al., 2013; Bar & Aminoff, 2003; Baumann & Mattingley, 2016; Gronau et al., 2008; Montaldi et al., 1998; Rémy et al., 2013; see also Aminoff, Gronau, & Bar, 2007) rather than - or at least in addition to - spatial layouts (Epstein, 2008; Epstein & Ward, 2009; Mullally & Maguire, 2011). Indeed, the PHC showed greater activity for incongruent vs. congruent images, though all scenes were identical in spatial layout, aside from the congruent/incongruent object that was pasted onto them. Furthermore, our study overcomes another, more specific possible confound (see Rémy et al., 2013, for discussion), which relates to spatial frequencies. The latter were found to modulate PHC activity (Andrews, Clarke, Pell, & Hartley, 2010; Rajimehr, Devaney, Bilenko, Young, & Tootell, 2011), implying that its object-related activation might be at least partially explained by different spatial properties of these stimuli, rather than by their semantic content. In the current study, all scenes were empirically tested for low-level visual differences between the conditions, including spatial frequencies, and no systematic difference was found between congruent and incongruent images (for details, see Mudrik et al., 2010), suggesting that the scenes differ mainly - if not exclusively - in contextual relations between the object and the scene in which it appears. Thus, our results go against low-level accounts of PHC activity, and suggest that it more likely reflects associative processes. This is in line with previous findings of PHC activations in response to objects which entail strong, as opposed to weak, contextual associations (Bar & Aminoff, 2003; Kveraga et al., 2011).

Finally, our results strengthen the claim that the prefrontal cortex (PFC; more specifically here – the inferior frontal gyri (see also Gronau et al., 2008; Rémy et al., 2013) and the cingulate cortex (see again Gronau et al., 2008)) is involved in object-scene integration per se. Some have suggested that frontal activations recorded during the processing of incongruent scenes are task-dependent (Rémy et al., 2013). Arguably, these activations arise since subjects need to inhibit the competing response induced by the scene’s gist (indeed, PFC activations - mostly in the right hemisphere - are found in response inhibition tasks; see Aron, Robbins, & Poldrack, 2004; Garavan, Ross, Murphy, Roche, & Stein, 2002; Garavan, Ross, & Stein, 1999). Yet in our experiment, subjects were not asked to give any content-related responses, but simply to freely view the stimulus sequence and indicate how visible it was. In this passive viewing condition, we still found PFC activations - mainly in the inferior and middle frontal gyri - bilaterally, while response inhibition is held to be mediated by the right PFC (Garavan et al., 1999). Our results accord with previous studies which implicated the PFC in associative processing, for related and unrelated objects (Gronau et al., 2008), for objects which have strong contextual associations (Kveraga et al., 2011), for related and unrelated words (Gold et al., 2006) and for sentence endings that were either unrelated (e.g., Baumgaertner, Weiller, & Büchel, 2002; Kiehl, Laurens, & Liddle, 2002) or defied world-knowledge expectations (Hagoort, Hald, Bastiaansen, & Petersson, 2004). In addition, increased mPFC connectivity with sensory areas has been found for congruent scenes which were better remembered than incongruent ones (van Kesteren et al., 2010), suggesting that this area mediates contextual facilitation of congruent object processing, and may be involved in extracting regularities across episodic experiences (Kroes & Fernández, 2012; Schlichting & Preston, 2015).

Taken together, our findings seem to support the model suggested by Bar (2004) for scene processing. According to this model, during normal scene perception the visual cortex projects a blurred, low spatial-frequency representation early and rapidly to the prefrontal cortex (PFC) and parahippocampal cortex (PHC). This projection is considerably faster than the detailed parvocellular analysis, and presumably takes place in the magnocellular pathway (Graboi & Lisman, 2003; Merigan & Maunsell, 1993). In the PHC, this coarse information activates an experience-based prediction about the scene’s context or its gist. Indeed, the gist of visual scenes can be extracted even with very short presentation durations of 13ms (Potter, Wyble, Hagmann, & McCourt, 2014; see also Joubert et al., 2005, though see Maguire & Howe, 2016), and such fast processing is held to be based on global analysis of low-level features (Malcolm et al., 2016). Then, this schema is projected to the inferior temporal cortex (ITC), where a set of schema-congruent representations is activated so that representations of objects that are more strongly related to the schema are more activated, hereby facilitating their future identification. In parallel, the upcoming visual information of the target object selected by foveal vision and attention activates information in the PFC that subsequently sensitizes the most likely candidate interpretations of that individual object, irrespective of context (Bar & Aminoff, 2003). In the ITC, the intersection between the schema-congruent representations and the candidate interpretations of the target object results in the reliable selection of a single identity. For example, if the most likely object representations in PFC include a computer, a television set, and a microwave, and the most likely contextual representation in the PHC correspond to informatics, the computer alternative is selected in the ITC, and all other candidates are suppressed. Then, further inspection allows refinement of this identity (for example, from a “computer" to a "laptop").

Based on this model, we can now speculate about the mechanisms of incongruent scenes processing: in this case, the process should be prolonged and require additional analysis - as implied by the elevated activations in all three areas in our study in the incongruent condition. There, since the visual properties of the incongruent object generate different guesses about object-identities in the PFC than the schema-congruent representations subsequently activated in the ITC, the intersection fails to yield an identification of the object, requiring further inspection and a re-evaluation of both the extracted gist (PHC) and the possible object-guesses (PFC). This suggestion can explain the widely-reported disadvantage in identifying incongruent objects, both with respect to accuracy (e.g., Biederman et al., 1974; Boyce et al., 1989; Underwood, 2005) and to reaction times (e.g., Davenport & Potter, 2004; Palmer, 1975; Rieger et al., 2008). It is further strengthened by EEG findings, showing that the waveforms induced by congruent and incongruent scenes start to diverge in the N300 time window (200-300 ms after the scene had been presented) - if not earlier (Guillaume, Tinard, Baier, & Dufau, 2016), at which these matching procedures presumably take place, prior to object identification (Mudrik et al., 2010; Mudrik, Lamy, et al., 2014; Võ & Wolfe, 2013; though see Ganis & Kutas, 2003 and Demiral et al., 2012). Note however, that this model holds the ITC as the locus of object-context integration, at which the information from the PHC and PFC converge. It also focuses on the perceptual aspect of object-context integration, to explain how scene gist affects object identification. Yet PFC activations (more specifically, IFG activations) were found also for verbal stimuli that were either semantically anomalous or defied world-knowledge expectations (for review, see Hagoort, Baggio, & Willems, 2009), suggesting that (a) the PFC may also be involved in the integrative process itself, rather than only in generating possible guesses about object identities irrespective of context and (b) that it may mediate a more general, amodal mechanism of integration and evaluation.

### 4.2. The role of consciousness in object-context integration

In this study, we found no neural evidence for unconscious object-context integration: using only trials in which subjects reported seeing nothing or only a glimpse of the stimulus, and were at chance in discriminating target image congruency (though slightly above chance in discriminating target image orientation), the BOLD activations found in the visible and localizer conditions became undetectable in our setup. Following previous studies which failed to find univariate effects during unconscious processing but managed to show significant decoding (e.g., Sterzer, Haynes, & Rees, 2008) we used multivariate pattern analysis (MVPA; Norman, Polyn, Detre, & Haxby, 2006) as a more sensitive way to detect neural activations which may subserve unconscious object-context integration; however, we did not find significant decoding in the invisible condition.

How should this null result be interpreted? One possibility is that it reflects the brain’s inability to process the relations between an object and a scene when both are invisible. This finding goes against our original behavioral finding of differential suppression durations for congruent vs. incongruent scenes (Mudrik, Breska, et al., 2011; see also Mudrik & Koch, 2013); however, a recent study failed to replicate (Moors et al., 2016) this original finding, and we too were not able to reproduce behavioral evidence for unconscious scene-object integration (Kataev & Mudrik, under review). Furthermore, another study which focused on the processing of implied motion in real-life scenes also failed to find evidence of unconscious processing (Faivre & Koch, 2014). In the same vein, the findings of another study which investigated high-level contextual unconscious integration of words into congruent and incongruent sentences (Sklar et al., 2012) was recently criticized (Shanks, 2016; for a review of studies showing different types of unconscious integration, see Mudrik et al., 2014). Taken together with the null result in our study, this could imply that high-level integration is actually not possible in the absence of awareness. This interpretation is in line with the prominent theories of consciousness which consider consciousness and integration to be intimately related (Dehaene & Changeux, 2011; Dehaene & Naccache, 2001), if not equivalent to each other (Tononi et al., 2016; Tononi & Edelman, 1998).

On the other hand, one should be cautious in interpreting the absence of evidence as evidence of absence. The observable correlates of unconscious integration processes may be so weak that we missed them; our study was likely underpowered both with respect to number of subjects and number of trials, and fMRI may simply not be a sensitive enough methodology to detect the weak effects of unconscious higher-level processing. The Bayesian analyses we performed suggest that our data in the invisible condition was inconclusive, and did not support the null hypothesis. Many previous fMRI studies either showed substantially reduced or no activations to invisible stimuli (Bahmani, Murayama, Logothetis, & Keliris, 2014; Hesselmann & Malach, 2011; Yuval-Greenberg & Heeger, 2013), or effects that were significant, yet weaker and more focused compared with conscious processing (e.g., Dehaene et al., 2001; Sadaghiani et al., 2010; Van Gaal, Lamme, Fahrenfort, & Ridderinkhof, 2011). Together, this raises the possibility that in some cases behavioral measures may be more sensitive than imaging results; in a recent study, for example, invisible reward cues improved subjects’ performance to the same extent as visible ones - yet while the latter evoked activations in several brain regions (namely, motor and premotor cortex and inferior parietal lobe), the former did not (Bijleveld et al., 2014). Critically, in that study subjects were performing a task while scanned. Our study, on the other hand, included no task in order to make sure that the observed activations were not task-induced, but rather represented object-context integration per se. Thus, while our findings cannot rule out the possibility that object-context integration indeed does not depend on conscious perception; they surely do not support this claim.

## Conclusions

While finding no definite evidence for (or against) unconscious object-context integration, the present study contributes to our understanding of conscious object-context integration. We found enhanced LOC, ITC, PHC and PFC activations for incongruent scenes, irrespective of task requirements. Our results cannot be explained by low-level differences between the images, including spatial frequencies, which were suggested as a possible confound in previous studies (Rémy et al., 2013). The use of stimuli that depict people performing real-life actions with different objects, rather than non-ecological stimuli (e.g., line drawings; Kim & Biederman, 2010; isolated, floating objects; Gronau et al., 2008; Kaiser et al., 2014), or stimuli with which subjects have less hands-on, everyday experience (animals or objects presented in natural vs. urban sceneries; Jenkins et al., 2010; Kirk, 2008; Rémy et al., 2013), enables us to better track real-life object-context integration processes which also occur outside the laboratory. Our results thus imply that in such real-life integration processes, incoming visual information about object features is compared with scene-congruent representations evoked by the scene gist, in an interplay between the abovementioned areas. This interplay - usually leading to contextual facilitation of object processing - is disrupted when incongruent scenes are presented, resulting in additional neural processing to resolve these incongruencies.

## Acknowledgments

We thank Christof Koch and Ralph Adolphs for constructive suggestions and helpful discussion, and Hagar Gelbard Sagiv and Uri Maoz for their help with the design. We further thank Galit Yovel and Nurit Gronau for their insightful comments on the manuscript. LM was supported by the Israel Science Foundation (grant No. 1847/16) and the Marie Sklodowska-Curie Individual Fellowships (grant No. 659765- MSCA-IF-EF-ST). NF was supported by the Fyssen and the Philippe foundations.

## Supplementary material

**Supplementary figure 1.**
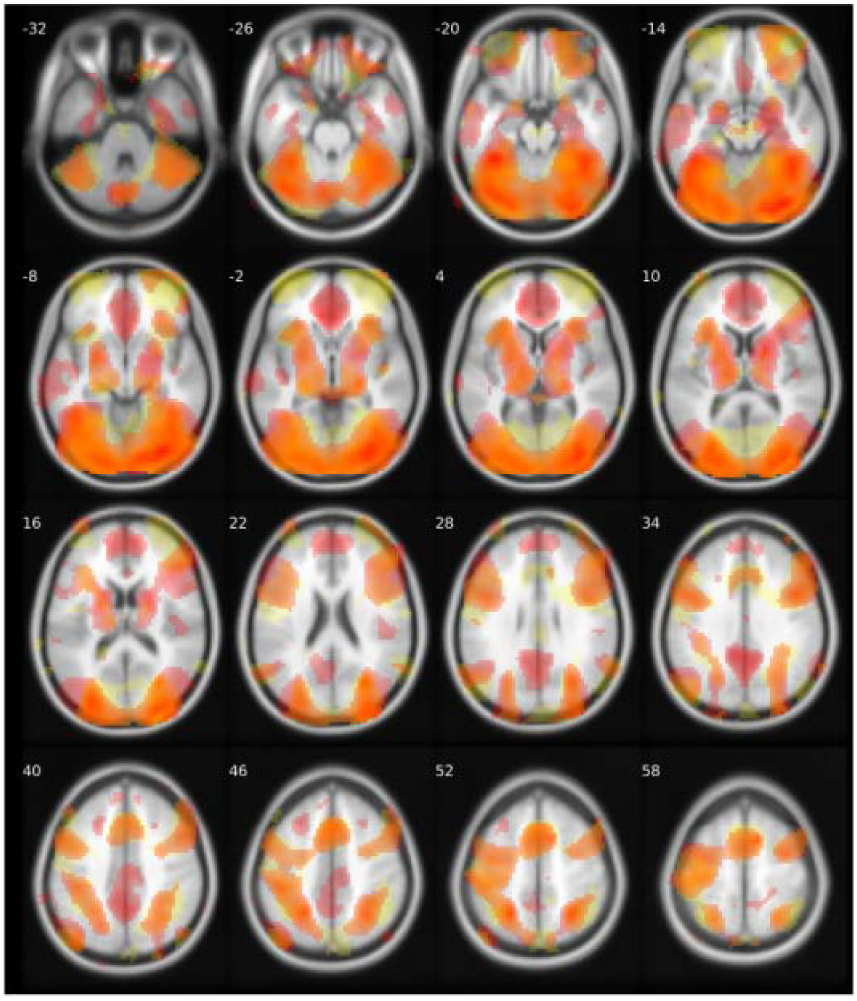
Masks of t-maps for the comparison of baseline vs. stimulation (congruent and incongruent) trials in the visible condition (in red), and invisible condition (in yellow). p < 0.001, uncorrected.

